# Computer-aided drug design and pharmacophore modeling towards the discovery of novel anti-Ebola agents

**DOI:** 10.1101/2025.03.23.644818

**Authors:** Io Diakou, George P. Chrousos, Elias Eliopoulos, Dimitrios Vlachakis

**Author notes:** Correspondence to*: Professor Dimitrios Vlachakis, Laboratory of Genetics, Department of Biotechnology, School of Applied Biology and Biotechnology, Agricultural University of Athens, 75 Iera Odos, 11855 Athens, Greece E⍰mail.

## Abstract

The Ebola virus belongs to the *Filoviridae* family of negative-sense RNA viruses and has been responsible for multiple widespread and severe outbreaks of a type of hemorrhagic fever, termed Ebola virus disease, mainly in the African continent. The virus, which is suspected to have a zoonotic origin, constitutes one of the most lethal pathogens, with high transmission rates and lasting effects on survivors. Research on the Ebola virus is made challenging due to a need for strict biosafety measures, while the arsenal of prophylactic and therapeutic measures remains limited. Modern computational methods can accelerate the process of developing candidate therapeutics that target essential proteins for the Ebola virus. In the present study, a pipeline was constructed to employ machine learning, big chemical data and structural bioinformatics towards the generation of novel molecules and the design of pharmacophores targeting the Ebola virus polymerase.

## Introduction

The *Filoviridae* family of RNA viruses includes the highly contagious and lethal Ebola virus, which was initially identified in 1976 in Sudan and the Democratic Republic of the Congo (formerly Zaire), where it was responsible for two epidemics of severe hemorrhagic fever that occurred at the same time (1). The first-recorded epidemic of the disease caused by the virus, termed Ebola virus disease (EVD), occurred close to the Ebola River; thus, the virus was named Ebola. Up to 90% case fatality rates have been observed during certain outbreaks of the virus, which is regarded as one of the most lethal viral pathogens worldwide (2). The Zaire, Sudan, Bundibugyo, Reston, and Tai Forest ebolaviruses constitute the five species of Ebola viruses that are currently known to exist (3). A linear, negative-sense RNA genome is contained in filamentous, pleomorphic viral particles that encase the encapsulated Ebola virus (4).

On January 11, 2023, the most recent Ebola virus illness outbreak, was declared. According to the World Health Organization (WHO), the epidemic began in September, 2022 in Mubende District, Uganda, and was the fifth known outbreak caused by the Sudan ebolavirus species. There were 142 confirmed cases, 55 fatalities and 22 suspected cases in the epidemic (5). The West African Ebola epidemic, which began on March 23, 2014, in Guinea and was caused by the Zaire ebolavirus species, was the largest Ebola outbreak in history. In terms of worldwide public health, that epidemic resulted in ∼28,652 total cases, 11,325 fatalities and a mortality rate of 40-70% (6). Guinea, Liberia and Sierra Leone were countries that were mostly affected by the epidemic; as these countries were already afflicted by war and poverty, the provision of essential care and containment measures was difficult (7). Aside from the casualties themselves, the outbreak had an impact on other healthcare services, such as maternity and child health, the treatment of human immunodeficiency virus (HIV) and running vaccine programs (8). Ebolaviruses are considered to be zoonotic viruses, which means they are suspected to be transferred from an animal to a human via direct contact with bodily fluids from infected animals, or through contact with items contaminated by these bodily fluids. Humans, once infected, can further transmit the illness by coming into close touch with body fluids, such as blood or saliva, or contaminated items and surfaces (9). A spillover event occurs when a disease, such as a virus, spreads from one species to another species that is not its reservoir host (10). Understanding spillover occurrences is critical for preventing the spread of emerging infectious illnesses. It is considered that three species of fruit bats of the *Pteropodidae* family are the natural reservoir host of the Zaire ebolavirus, following various modeling and serological experiments (11). The Ebola virus enters the human body through mucous membranes or skin breaches. Once in the body, the virus infects immune cells, particularly macrophages and dendritic cells, and other cell types such as liver cells, endothelial cells and fibroblasts (12). The incubation period for EVD can range from 2 to 21 days, with an average of 5 to 7 days. The virus replicates in the body of the host throughout this asymptomatic incubation phase, and the host can only transmit the infection when symptoms appear (13). Typical symptoms of EVD in the early stages include fever, headache, jaundice, myalgia and gastrointestinal symptoms such as nausea, vomiting and diarrhea (13).. Infected patients may experience complications such as internal bleeding or multi-organ malfunction as the illness advances (13) . For diagnosis, the virus can be detected using various techniques, such as enzyme-linked immunosorbent assay (ELISA), reverse transcription-polymerase chain reaction (RT-PCR), serological tests and on-field rapid diagnostic kits (14). The handling of patients and patient samples requires the use of personal protective equipment by medical personnel and stringent decontamination regimes, while the study of viral samples requires biosafety level 4-grade facilities (15).

The Ebola virus particle or virion is a complex structure comprised of multiple unique components. The single-stranded RNA genome, which runs ∼19,000 base pairs in length, is at the heart of the virion. This genome is encircled by viral structural proteins, which join together to form the inner ribonucleoprotein (RNP) complex (16). The nucleoprotein (NP) and the RNA-dependent RNA polymerase (L) are two key components of the RNP complex. These proteins perform critical functions in viral replication, ensuring that the virus can reproduce and propagate inside the host cells (17). The viral envelope, which is comprised of two viral glycoproteins, GP1 and GP2, surrounds the RNP complex. The glycoproteins interact with host cell surface receptors, such as C-type lectin family receptors, to enable attachment of the virus and membrane fusion for subsequent entrance into cells (18). Knowledge of the Ebola virus structure and replication cycle provides a foundation for the development of prophylactic and therapeutic strategies. Notable approved anti-Ebola vaccines include rVSV-ZEBOV-GP (ERVEBO), Ad5-EBOV and rVSV/Ad5, all expressing the ebolavirus glycoprotein as the immune system-stimulating antigen (19–21). Treatment strategies encompass various types, such as the convalescent serum of infected individuals, therapeutic interferons, gene therapies such as small interfering RNAs that target viral genes and lastly, inhibitors and monoclonal antibodies (22). Inhibitors may target various host machinery or viral enzymes crucial for the ebolavirus life cycle; ion channel inhibitors target cell ion channels to inhibit viral entry and viral entry inhibitors target host or viral enzymes crucial for viral attachment, viral fusion and entry (23). Moreover, monoclonal antibodies have been approved for the treatment of EVD; Inmazeb and the more recent Ebanga both target the ebolavirus glycoprotein, thus inhibiting viral entry and attenuating viral spread across host cells (24,25). Lastly, nucleotide analogues, such as Remdesivir, become incorporated into nascent RNA molecules during the viral RNA synthesis and thus interfere with viral replication (26).

The threat of Ebola virus remains, as demonstrated by the recurrent outbreaks even well into 2023, reaffirming the need for safe and effective prophylactic measures and treatments. Research on the Ebola virus is made challenging by the requirement for austere safety measures in the laboratory setting, to prevent accidental transmission of the pathogen (17). Furthermore, the drug development process itself is a time-consuming and costly process from target identification to lead development and optimization, with a fraction of drugs under development finally making it to market (27). Data abundance marks the current state of scientific knowledge; a trove of structural, genetic, chemical and molecular data is harnessed and is available in public databases (28,29). Machine learning and deep learning solutions, additionally, are being increasingly used in the context of the drug development process (30). A risk prediction model for lymph node metastasis in Ewing’s sarcoma (ES) has been created using clinicopathological data leading to the design of a clinical-use service to improve detection of this metastasis in patients with ES (31). Similarly, a model was trained with medical data from patients with kidney cancer to aid in the stratification for the risk of developing bone metastasis and inform personalized treatment decisions (32).

Emerging technologies, such as semantic analysis or neural networks can be employed in tasks such as target identification or drug development against single-stranded RNA viruses (28). The use of computational methods has the potential to accelerate the preclinical drug development of novel antivirals, reducing costs and removing the need for biosafety measures. The present study describes a computational pipeline for the generation of new molecules targeting the Ebola virus polymerase, employing neural network architectures, chemical data and structural bioinformatics. The use of public big data of existing anti-virals enables the training of models on critical features of anti-viral agents, which in turns informs the design of novel candidates by the neural network. The *in silico* assessment of the neural network designed ligands by computational bioinformatics and the design of pharmacophores further compresses these features and creates a profile of antiviral potential, capable of informing further experiments. The flexibility and scalability of the pipeline components enhances and accelerates research on the preclinical stage, where scale and safety are all the more relevant when faced with the challenge of viral epidemics and pandemics.

## Data and methods

### Construction of pipeline

The overview of the pipeline is summarized in Fig. 1A and is as follows: Public chemical databases were employed towards building an in-house chemical database of existing compounds which target viral polymerases. A neural network was designed and trained on the in-house database to learn the underlying features and distribution. Subsequently, the latent space defined by the network was explored to generate new molecules with a set of desired properties. The resulting novel ligands were further filtered and a conformational search was subsequently carried out to generate three-dimensional conformations of the molecules, in preparation for a docking simulation. A high throughput screening protocol was used to dock the ligands against the previously experimentally determined, three-dimensional structure of the Ebola virus polymerase-VP35 complex, and lead performing ligands served as the basis for pharmacophore construction.

**Figure 1.**
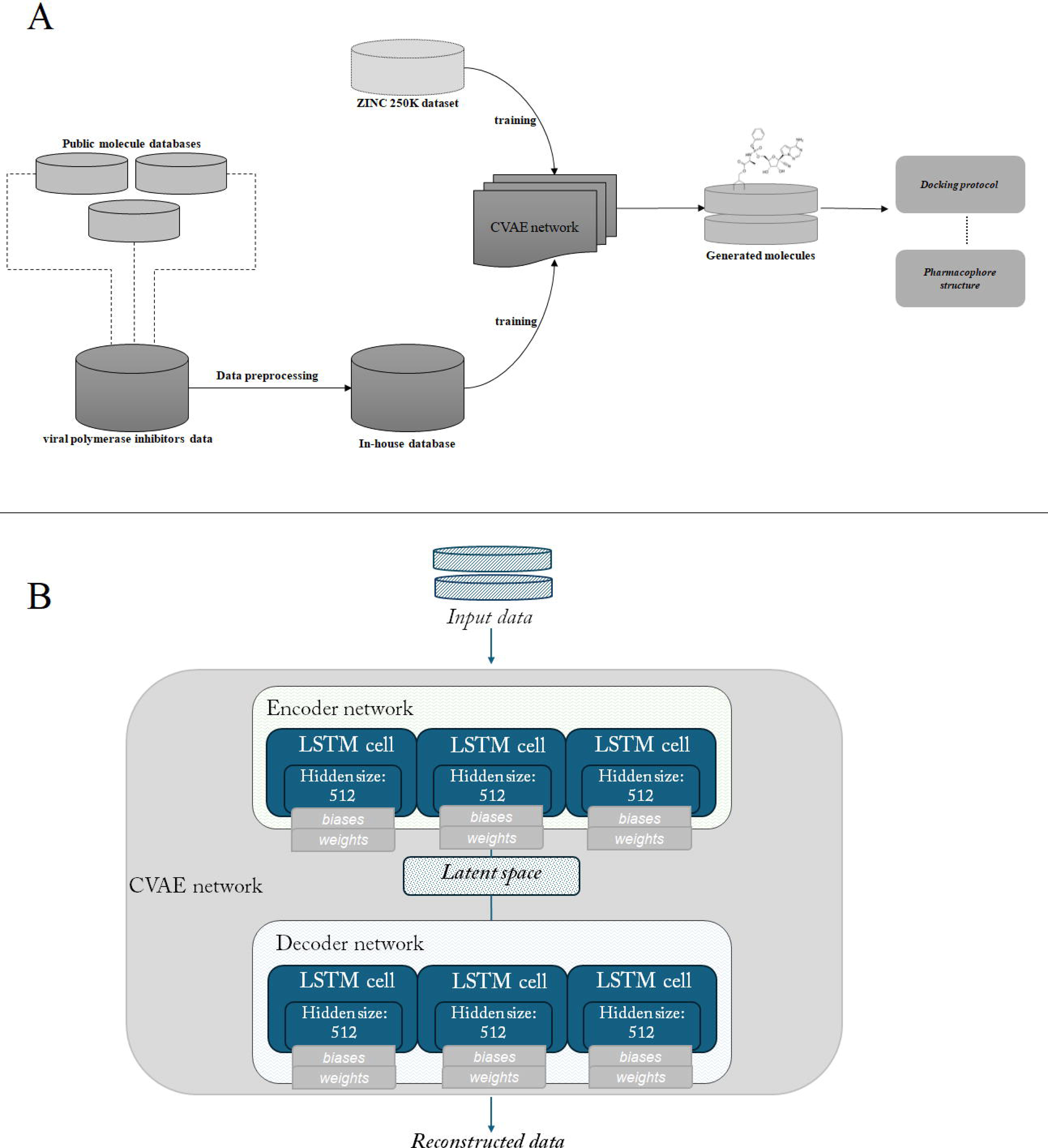
Overview of the pipeline and the network architecture. (A) Schematic view of the pipeline steps. (B) Schematic view of the variational autoencoder’s architecture and cells. CVAE, conditional variational autoencoder; LSTM cell, long short-term memory cell.

### Creation of in-house chemical database and data pre-processing

In the era of big data and high-throughput processes, vast amounts of chemical data are available in openly accessible databases. ChEMBL constitutes a widely-known, public database of curated chemicals, containing more than two million preclinical drugs with associated bioactivities, ∼14,000 clinical candidate drugs and >4,000 clinical drugs (33). Its sister database, SureChEMBL, aggregates similar data (34). The Kyoto Encyclopedia of Genes and Genomes (KEGG) Drug database collects drug data across three different continents in a systematic manner (35). The Virus Pathogen Database and Analysis Resource (ViPR) hosts multi-omics data regarding various viral families, combining data from archives, as well as data submitted by scientific groups and virologists, rendering it a valuable source of information (36). PubChem, supported by the National Library of Medicine of NIH, constitutes a principal information source for medicinal chemistry and drug discovery, with substance, compound and bioassay records available along with various metadata (37).

Modules were built in Python to use the Application Programming Interface (API) of ChEMBL, PubChem, KEGG and SureChEMBL to collect data of compounds which target viral polymerases. The selection of the data sources was steered by a set of defined criteria, such as the scope and coverage of the data, the public of private nature of the datasets, the format of the data, as well the availability of an API to enable the efficient sourcing. The chemical data were collected in simplified molecular input line entry system (SMILES) format. SMILES is used to convert the three-dimensional structure of a chemical’s into a string of symbols that can be used as input in neural network training tasks or other bioinformatic analyses (38). In the case of ViPR, where the API availability is not straightforward, the collection of compounds was executed in a semi-manual manner. The ChEMBL Python module, for instance, queries the ChEMBL molecule endpoint against a set of ChEMBL target identifiers of interest (such as the influenza polymerase, CHEMBL4523676) to collect compounds against that target, along with metadata such as compound preferred names, max phase, molecular formula, molecular weight and the canonical SMILES string. The module can be found in the following repository: https://github.com/IoDiakou/Automatic-collection-of-drugs-for-specific-target-from-ChEMBL. Analogous modules were constructed for all the aforementioned databases and the collections of compounds queried from each database were finally collected into an in-house SQLite database of viral inhibitors. SQLite offers flexible and scalable storage, which enables the preprocessing and handling of the data (39).

Upon establishing the database, preprocessing was carried out using the Python RDKit interface, which allows the easy manipulation and transformation of chemical strings (40). Functions were used to filter noisy compounds out of the in-house database; examples include mixtures, non-organic salts, organometallic compounds, polymers. Considerably lengthy molecules were also filtered out, as lengthy strings which exceed a threshold of ∼120 characters can hinder deep learning and cheminformatics tasks. Furthermore, to handle the existence of duplicate records, as a direct result of sourcing inhibitors from public databases where overlap may occur, a deduplication step was also carried out. Lastly, molecular properties of interest were calculated for each entry of the filtered and deduplicated compounds database.

### Neural network architecture design and training

Generative probabilistic models based on neural networks have gained traction over the past decade, with generative adversarial nets and variational autoencoders (VAEs) as two prominent examples, being applied to various tasks (41). The neural network learns a distribution p(x) across data points and allows sampling, thus carrying out the generative portion (42). The architecture used herein is a subtype of VAE, termed a conditional variational autoencoder (CVAE); the model samples from the generative distribution upon a set condition, to generate novel data points and simultaneously predict properties. All data handling and machine learning tasks were carried out using Python; the neural network model design and deployment was carried out using the TensorFlow framework, which offers scalability and reliability for machine-learning tasks (43). The VAE consists of two units, the encoder, which maps input data points to a distribution over the low-dimensional, latent space, and the decoder, which maps the points of the aforementioned latent space back into the data space. During training, the loss function implemented aims to balance two objectives. The reconstruction loss reflects how well the decoded data points match their original inputs and is calculated by the mean squared error (44). The regularization part of the loss function encourages the high similarity between the distribution which the neural network learns, and an a priori distribution, commonly a standard Gaussian distribution.

The encoder component of the network comprises of a list of Long short-term memory (LSTM) cells, with each LSTM cell in the list having a specific hidden size, and all LSTM cells are wrapped into a multi-layered LSTM cell. An analogous structure of LSTM cells is defined for the decoder. LSTMs belong to the umbrella group of recurrent neural networks and constitute, one might say, an evolution of them (45). A schematic of the configuration and cell structure of the neural network is presented in Fig. 1B. The designed neural network was trained sequentially on the ZINC 250K dataset and then the in-house database of viral polymerase inhibitors, to learn the representations and the characteristics underlining these viral inhibitors. The ZINC database and established subsets of it are commonly used in docking experiments or other string-based chemical research approaches; the 250K subset of the ZINC database has been used in Kaggle applications and other autoencoder-related articles (46).

### Generation and filtering of novel molecules

Upon being fine-tuned on the dataset of existing viral polymerase inhibitors, the CVAE was tasked with generating novel molecules in SMILES format. The generation was steered towards molecules which would exhibit the properties within the ‘neighborhood’ of an established small-molecule viral polymerase inhibitor. Both the training and the sampling stage were carried out in VS Code, using a system comprising of a quad-core Intel Core i7-3770 processor and an NVIDIA GeForce GXT 1650 architecture for GPU acceleration, which is crucial for machine-learning tasks.

The dataset of generated molecules was stored in a new SQLite table instance for the subsequent pipeline steps of re-filtering and docking experiments. RDKit functions and SMARTS patterns were used to flag compounds with high similarity to the starting in-house dataset, or with low likelihood of being synthesized or proceeding further during a conventional hit identification stage, such as fully acyclic compounds.

### Docking and pharmacophore generation

To investigate the potential efficacy of the novel molecules as inhibitors for the Ebola polymerase, docking experiments were carried out in molecular operating environment (MOE); docking is a popular technique in computational drug discovery for the study of ligand-target interactions (47). To transition from the generated molecules, which are in a 2-dimensional flat string format, to a three-dimensional molecule, conformational search protocols were carried out. The complex of EBOV polymerase and cofactor VP35 with suramin (PDB id: 7YET) served as the docking structure, following structure preparation steps of assigning formal charges, energy minimization and extraction of the suramin molecule in complex. The complexed suramin additionally served as a baseline for the binding site region as well as for interactions with surrounding residues and binding affinities. High-throughput screening constitutes a robust protocol for the rapid assay of candidate compounds, allowing for the time and computational resource-efficient evaluation of large sets of chemical data against a target (48). Pharmacophore modeling was carried out on the generated molecules with optimal binding scores and interactions; pharmacophores can serve as a framework for virtual screening efforts, allowing the identification of motifs and interactions of interest underlying activity (49).

## Results

### Construction of in-house chemical database

In the stage of database creation, API queries were built against public databases, such as PubChem and ViPR, for the search and retrieval of information related to molecules with inhibitory activity against viral polymerases. Prior to filtering, the in-house database of compounds contained 12,369 molecules; following deduplication and removing out-of-scope molecules, such as mixtures, non-organic salts and molecules >120 characters, the database contained 11,960 unique molecule entries. For each SMILES molecule, relevant molecular properties were calculated, namely molecular weight, logP and topological polar surface area (TPSA), serving as additional descriptors and features for the training of the machine-learning model. The TPSA represents the total of the contributions to the molecule (typically van der Waals) surface area of polar atoms, e.g., nitrogen, oxygen or associated hydrogens (50). It constitutes a popular molecular descriptor for tasks of evaluating transport ability, penetrative ability (such as through the blood-brain barrier) and absorption within the intestine (50).

### Neural network training

The SMILES input molecular data are one-hot encoded into their vector embedding, within the first layers of the encoding unit, along with the corresponding molecular properties previously calculated for each data entry. The latent representation is created in the encoder unit and subsequently serves as input for the decoder unit; the decoded vector generated by the decoder unit is funneled through a transformative layer of equal size to the original one-hot encoding vector of the encoding unit, to ensure the faithful reconstruction of the input. Softmax activation is applied to the vectors. To set the basis for the faithful reconstruction of encoded SMILES, the neural network was first trained for 10 epochs on the ZINC 250K dataset, using Adam as the optimizer, while also implementing an early-stopping criterion. Subsequently, the model was fine-tuned on the in-house database of existing inhibitors using ten-fold augmentation, to prepare for the generative step of the pipeline.

### Novel molecule generation, docking and pharmacophore design

The latent space constructed between the encoder and decoder unit served as the chemical space for the generation of novel molecules. The latent space was sampled for the generation of 1,400 novel molecules along with prediction of molecular properties for the generated molecules, resulting in sets of SMILES, properties. SMARTS patterns were employed, post-deduplication, for the removal of invalid SMILES strings from the resulting dataset. As a last step the generated molecules were evaluated for sufficient dissimilarity against the existing inhibitors within our in-house database, which served as the fine-tuning set.

To investigate the binding behavior and interactions between the generated molecules and the target of interest, herein the ebolavirus polymerase, the structure of the ebolavirus polymerase complex was first prepared in MOE in anticipation of the docking experiments. The employed structure consists of the Ebola virus L polymerase complexed with the co-factor VP35, an accessory protein that is an essential part of the replication unit, with supplementary roles as an antagonist of the host immune response and virulence factor (51). VP35, which is illustrated by the dark blue ribbon formation in Fig. 2A, interacts with the Ebola virus L protein (Fig. 2A with red, yellow and cyan ribbons for the different polymerase domains) to form the replication complex for the generation of viral genome strands during the viral cycle. In the employed three-dimensional structure (PDB ID: 7YET), suramin is additionally bound to the complex (52) (Fig. 2 in pink sphere representation to denote the span of its molecular surface).

**Figure 2.**
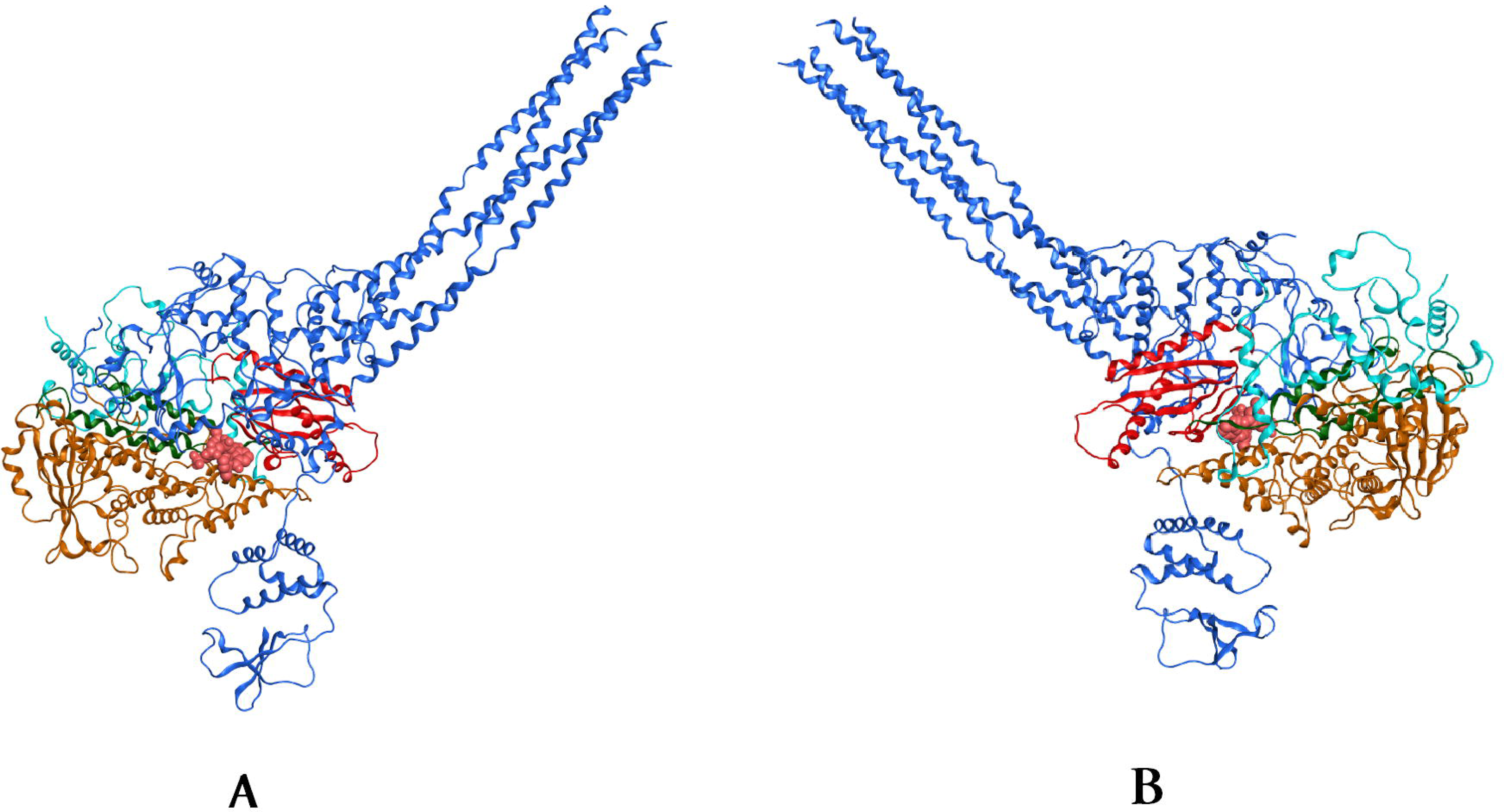
Complex of the Ebola virus L polymerase - VP35 cofactor and suramin. (A) Back view. (B) Front view (PDB ID: 7YET, image generated in MOE). The VP35 cofactor is colored in dark blue ribbon representation, the suramin molecule in the center in salmon pink spheres representation and the L polymerase in yellow, green, light blue and red ribbon representation.

The generated molecules underwent a protocol of conformational search within MOE to build a library of three-dimensional conformations, which are crucial for the exploration of the structural space of the binding site during the docking experiments. Detailed conformational search cycles were performed, with 100 iterations threshold per molecule for each conformation, and a set of ten conformations per molecule within the dataset. The suramin molecule which is bound to the active site of the Ebola virus L-VP35 complex served as a guideline for the identification of the site for docking, using dummy atoms to define the site and the receptor and solvent as the receptor parameter. The ligand database containing the conformations of the novel molecules was then docked against the site (53).

The docking results were collected in a MOE molecular database (mdb) file for further analysis, which encompassed the evaluation of the results on multiple levels. The docking scores and position of suramin within the site, upon using the same docking protocol and parameters, served as a baseline for the comparison of the scoring results of the novel molecules as well as for visual comparisons. Examples of docking positions of the novel molecules are illustrated in Fig. 3, with the positioning of suramin (pale pink coloring) for visual comparison. Energy minimization was additionally carried out to optimize the poses, monitoring the shift in the positioning of the docked conformer. This is exemplified in Fig. 3, where the positioning of the ligand (light blue) relative to the binding pocket and position of suramin (in pale pink) has shifted following energy minimization cycles.

**Figure 3.**
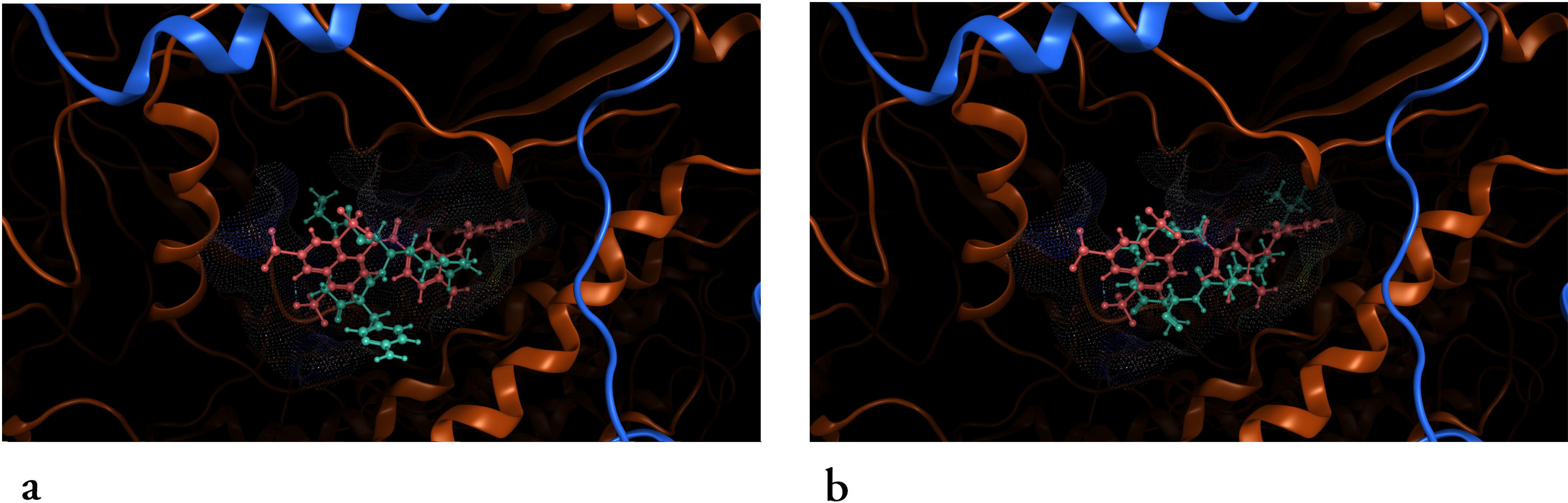
Conformations of a generated molecule (termed mol_2) and suramin, (A) before and (B) after energy minimization of the docked pose of mol_2. Suramin colored in salmon pink, mol_2 colored in light blue, both in ball-and-stick visualization form, captured in MOE. The molecular surface of the L-VP35 docking site in ‘dot’ visualization can be seen forming a confine around the two compounds.

Furthermore, the ligand interaction functionality within MOE allowed the examination of the specific interactions between the atoms of the molecules and crucial residues around the binding site of the viral polymerase. Conformational and spatial characteristics, such as steric hindrances, were also taken into account during the analysis of the docking results; structural characteristics, such as the absence or presence of highly cyclic groups, can serve as additional indicators of the molecules’ nature, such as their potential toxicity. Fig. 4 depicts this type of interactions information for the same docked molecule as in Fig. 3, termed mol_2 for brevity.

**Figure 4.**
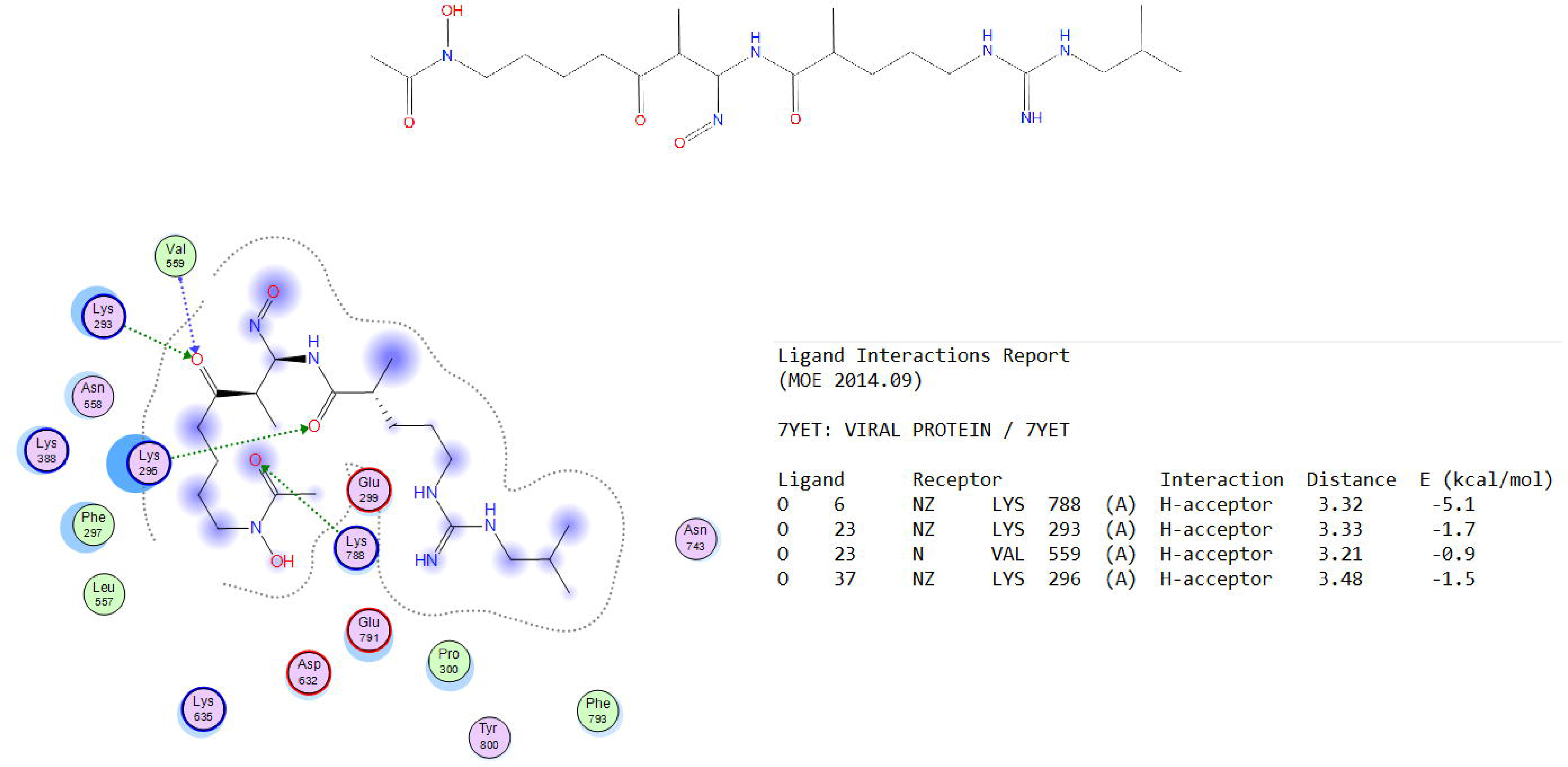
Details of docked molecule mol_2. (Top panel) SMILES sequence of the generated molecule. (Bottom left panel) Ligand interactions diagram generated in MOE. (Bottom right panel) Ligand interactions report generated in MOE. SMILES, simplified molecular input line entry system; MOE, molecular operating environment.

The interaction diagram illustrated in Fig. 4, on the bottom left panel, confirms that mol_2, for instance, forms several hydrogen bonds with the receptor ebolavirus protein complex. Key residues involved in these interactions are Lys 788, Lys 293, Val 559 and Lys 296. The hydrogen bonds, indicated by green dashed lines, suggest that these interactions are key for stabilizing the ligand-receptor complex. The interaction table provides quantitative details, with interaction energies ranging from -0.9 to -5.1 kcal/mol. These values indicate the strength of the hydrogen bonds, with the interaction between mol_2 atom O6 and EBOV polymerase Lys 788 being the most significant due to its lowest energy (-5.1 kcal/mol). The distances of these interactions (around 3.2-3.5 Å) are typical for hydrogen bonds, further validating the strength and specificity of these interactions. Top performing molecules as identified during this evaluation process were further optimized with molecular dynamics simulations without restraints, monitoring their shifts in positions as in the example of Fig. 3. Lastly, pharmacophore modeling was carried out via the pharmacophore building utility of MOE, calculating consensus features from the bound ligands wherever meaningful.

To carry out the union of consensus pharmacophores generated by different schemes, for each scheme, the individual pharmacophores were first generated per ligand member among the top 100 scoring docked ligands. The R-strength value was enabled, and each feature was adjusted with an appropriate assignment of weights, after assessing its importance in the set of overall interactions between the ligand and the receptor residues. Pharmacophore data were saved in the pharmacophore format of MOE, to be subsequently used to build the consensus pharmacophore for the scheme by combining the individual pharmacophore features from all ligands. As illustrated in Figs. 5 and 6, the unified pharmacophore features span the pocket defined by the docking site, reflecting interactions between the novel ligands and key residues of the viral enzyme. Overall, the binding site environment is comprised of a mix of hydrophobic and polar/charged residues. This diverse environment supports a range of interactions, from hydrophobic and aromatic interactions to hydrogen bonding and ionic interactions. Key residues of the receptor harboring potential interactions with features of the ligand are highlighted in yellow boxes in depicted Fig. 6, colored dark green and in stick figure representation as captured in MOE. Notably, Lys393, Glu398, Val583, Glu394 and Glu399 are highlighted, as these are key residues for binding interactions, highlighted by the analyzed interactions of suramin as well. Aromatic interactions, as highlighted in the generated pharmacophores, play a pivotal role in the ligand binding, particularly through π-π stacking interactions.

**Figure 5.**
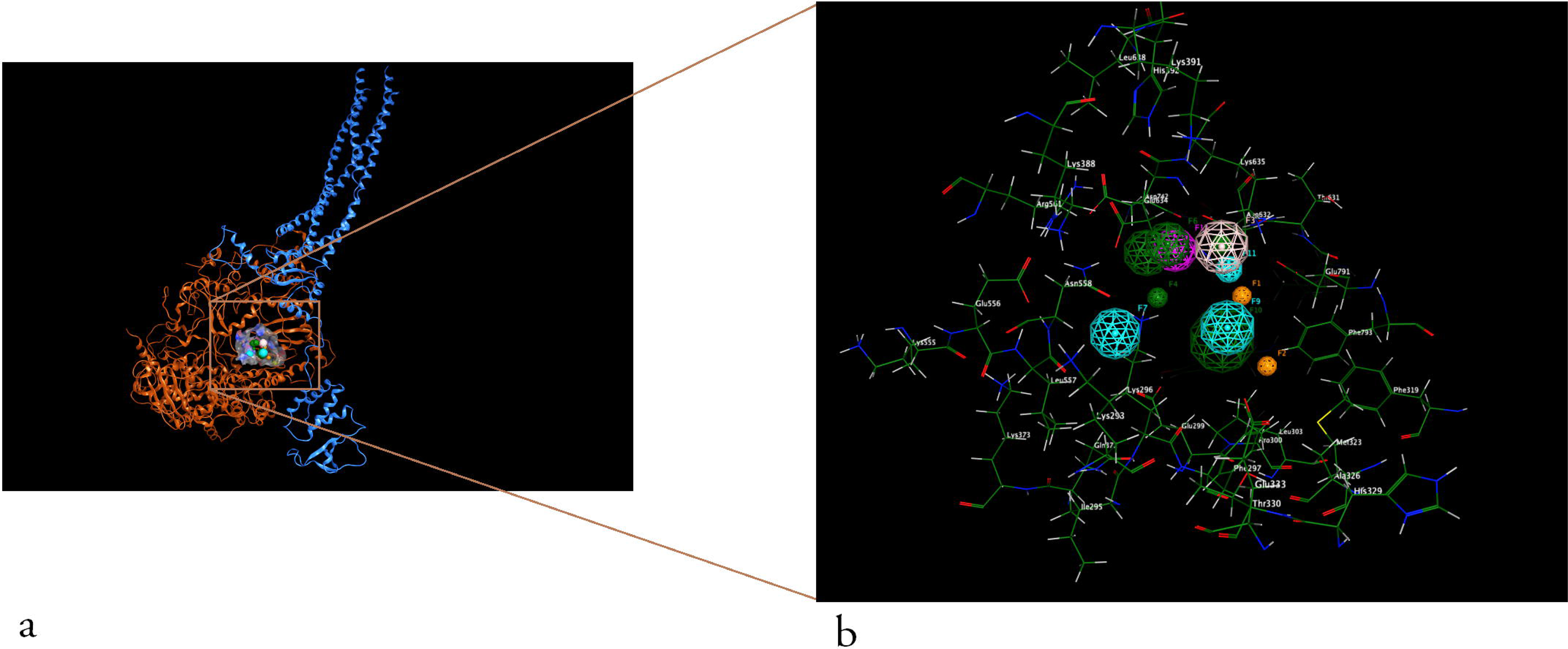
Consensus pharmacophore designed based on the top 100 docked ligands. (A) The pharmacophore position with regards to the EBOV VP35-L complex (back view). (B) Detailed view of the pharmacophore in the docking site with residues in dark green captured in MOE. MOE, molecular operating environment.

**Figure 6.**
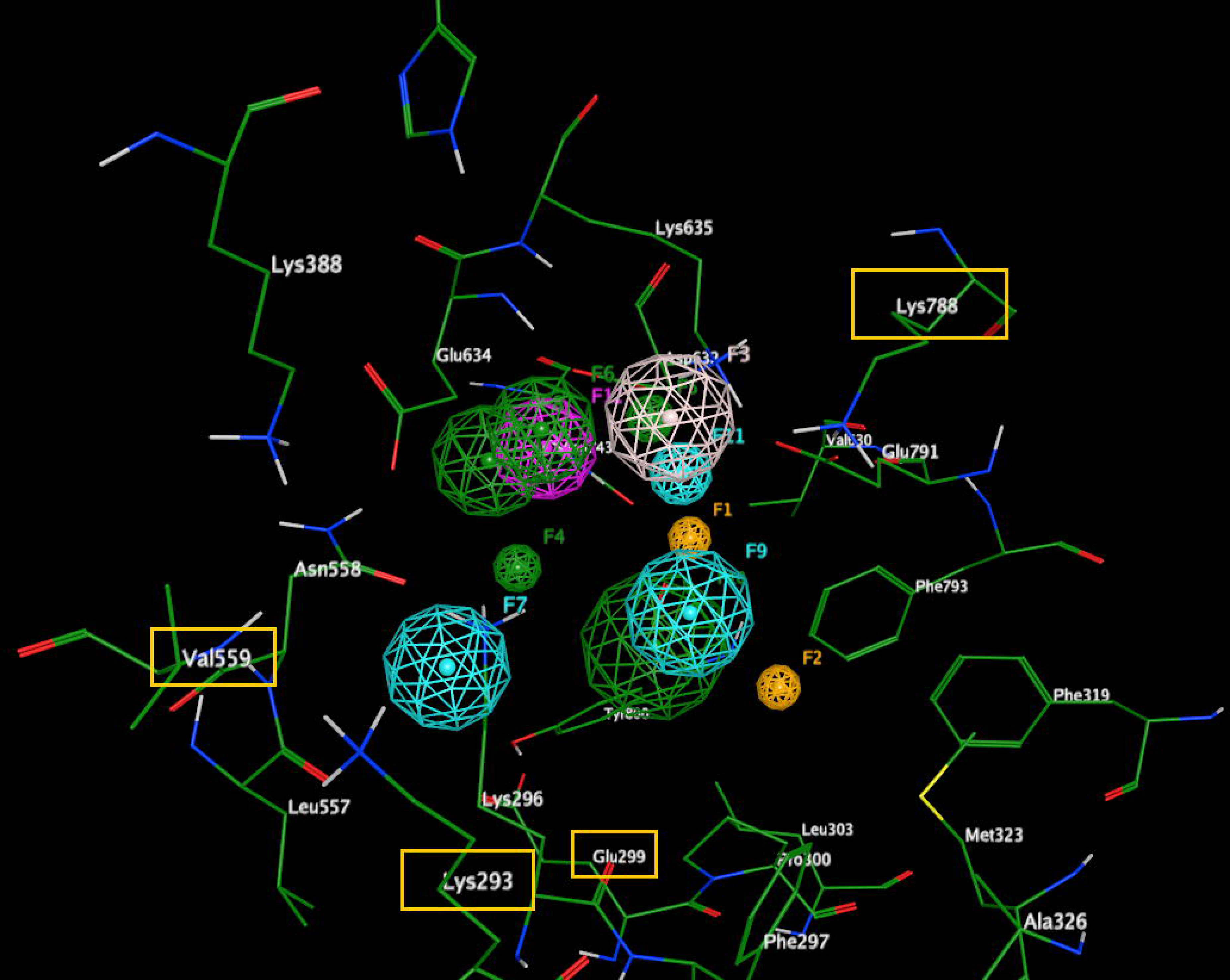
Highlights of critical receptor residues in the consensus pharmacophore. The figure summarizes specific Ebola virus polymerase residues, contained in the yellow boxes, harboring potential interactions with features of the ligand.

The unified pharmacophore reflects a comprehensive interaction profile, capable of forming multiple types of interactions within the receptor binding site, spanning aromatic, hydrophobic, hydrogen bonding and potential metal ion coordination interactions. Furthermore, this rich set of binding and chemical feature information can be employed in the subsequent design or screening of candidates against the specific target, employing the availed pharmacophoric features as quick guidelines for the design, screening or optimization of compounds and leads. Within the context of the pipeline described herein, the data which were collected during the docking and pharmacophore step of the pipeline were fed back into the SQLite database, linked as metadata to the SMILES molecules and further enhancing the knowledgebase.

## Discussion

The pipeline described herein constitutes a scalable approach for the discovery of novel potentially active anti-Ebola agents, harnessing the rapidly increasing power of machine learning capabilities and publicly accessible chemical data. Integral processes of the drug design challenge, such as the hit identification and lead optimization, can be supported by state-of-the-art computational methods, which have the potential of bringing vetted candidate molecules to the preclinical and laboratory stage, effectively reducing operational costs and time, both of which are crucial in the drug development cycle. This is all the more relevant when the target of interest is a virus, which, as exhibited by the recent COVID-19 epidemic, can grow into a global issue rapidly. Through the implementation of rapid and effective computational pipelines which combine string-based formats of drugs and active compounds, with the power of robust neural networks, a shift can be supported towards the adoption of a proactive stance in the development of safe and effective anti-virals.

Such a pipeline is marked by key features of scalability and repurposing. The training dataset and the target of interest are vessels, able to adopt different identities, depending on the task at hand. What is described herein as the design and evaluation of candidate, novel polymerase inhibitors targeting the EBOV polymerase can be efficiently translated into the generation of novel immunosuppressants in the context of Lupus, for instance, or the generation of novel inhibitors against a metabolic enzyme implicated in metabolic disorders. Furthermore, the neural network architecture itself constitutes a flexible parameter of great impact; tuning different parameters such as the network cell of choice or the optimization algorithm can in turn yield different generative behaviors, rendering the generation of novel data points, or molecules, more or less restrictive or more or less imaginative. One must note that there are inherent challenges in using SMILES as a representation of chemical information for the design and deployment of deep neural networks, including limited ability to reflect stereochemistry and challenges in encoding and interpreting complex molecules. Canonical SMILES can be employed to address the aspect of ambiguity, while the combination of SMILES with state-of-the-art tokenization techniques may enhance performance (54).

Processes similar to the framework described herein, such as the one proposed by van Tilborg and Grisoni (55), incorporate novel data acquisition to perform iterative model improvement, parallel to the model’s execution of the task, which, in the study by van Tilborg and Grisoni (55), is hit detection through screening. Despite not incorporating this factor of ‘active learning’, the proposed framework leverages the size and complexity of the training data of anti-virals that have been designed and assessed by the scientific and biomedical community, often in the lab setting. This allows the model to learn features from real-world examples, a ground truth of chemical information that leads to precise, multifaceted, in-depth learning and subsequent design of novel ligands. In summary, the proposed framework reflects the significant potential which exists in the application of machine-learning approaches in the field of computational drug design. Leveraging big data and state-of-the-art algorithms supports the effort of developing effective therapeutic agents, under the scope of precision medicine and a proactive stance against emerging viral threats.

## Acknowledgements

Not applicable.

## Availability of data and materials

The data generated in the present study may be requested from the corresponding author. The employed programming code and datasets can be accessed in the first author’s repository, in the following web link: https://github.com/IoDiakou/anti-ebola-drugs-vae.

## Authors’ contributions

ID, GPC, DV and EE have been involved in the design and conceptualization of this study. ID, GPC, DV and EE have contributed to the collection and analysis of the data, as well as the design and implementation of the pipeline described herein. ID, GPC, DV and EE have been involved in the writing and editing of the manuscript and the generation of relevant figures. All authors have read and approved the final manuscript.

## Ethics approval and consent to participate

Not applicable.

## Patient consent for publication

Not applicable.

## References

1. Clegg, J. and G. Lloyd, Ebola haemorrhagic fever in Zaire, 1995. Current Opinion in Infectious Diseases, 1995. 8: p. 225–228.

2. Muyembe-Tamfum, J.-J., et al., Ebola virus outbreaks in Africa: Past and Present. The Onderstepoort journal of veterinary research, 2012. 79: p. E1–E8.

3. Zheng, H., et al., Ebolavirus Classification Based on Natural Vectors. DNA and Cell Biology, 2015. 34(6): p. 418–428.

4. Ajelli, M., et al., Spatiotemporal dynamics of the Ebola epidemic in Guinea and implications for vaccination and disease elimination: a computational modeling analysis. BMC Medicine, 2016. 14(1): p. 130.

5. Branda, F., et al., The challenges of open data for future epidemic preparedness: The experience of the 2022 Ebolavirus outbreak in Uganda. Frontiers in Pharmacology, 2023. 14.

6. Kucharski, A.J. and W.J. Edmunds, Case fatality rate for Ebola virus disease in west Africa. The Lancet, 2014. 384(9950): p. 1260.

7. Gostin, L.O. and E.A. Friedman, A retrospective and prospective analysis of the west African Ebola virus disease epidemic: robust national health systems at the foundation and an empowered WHO at the apex. The Lancet, 2015. 385(9980): p. 1902–1909.

8. Brolin Ribacke, K.J., et al., Effects of the West Africa Ebola Virus Disease on Health-Care Utilization – A Systematic Review. Frontiers in Public Health, 2016. 4.

9. Marí Saéz, A., et al., Investigating the zoonotic origin of the West African Ebola epidemic. EMBO Molecular Medicine, 2015. 7(1): p. 17–23.

10. Plowright, R.K., et al., Pathways to zoonotic spillover. Nature Reviews Microbiology, 2017. 15(8): p. 502–510.

11. Koch, L.K., et al., Bats as putative Zaire ebolavirus reservoir hosts and their habitat suitability in Africa. Scientific Reports, 2020. 10(1): p. 14268.

12. Osterholm Michael, T., et al., Transmission of Ebola Viruses: What We Know and What We Do Not Know. mBio, 2015. 6(2): p. 10.1128/mbio.00137-15.

13. Velásquez, G.E., et al., Time From Infection to Disease and Infectiousness for Ebola Virus Disease, a Systematic Review. Clinical Infectious Diseases, 2015. 61(7): p. 1135–1140.

14. Bettini, A., D. Lapa, and A.R. Garbuglia, Diagnostics of Ebola virus. Frontiers in Public Health, 2023. 11.

15. Verbeek, J., et al., Personal protective equipment for preventing highly infectious diseases due to exposure to contaminated body fluids in healthcare staff. Cochrane Database of Systematic Reviews, 2020. 4.

16. Ghosh, S., et al., Genome structure and genetic diversity in the Ebola virus. Current Opinion in Pharmacology, 2021. 60: p. 83–90.

17. Judson, S., J. Prescott, and V. Munster *Understanding Ebola Virus Transmission*. Viruses, 2015. 7, 511–521 DOI: 10.3390/v7020511.

18. Nanbo, A., et al., Ebolavirus Is Internalized into Host Cells via Macropinocytosis in a Viral Glycoprotein-Dependent Manner. PLOS Pathogens, 2010. 6(9): p. e1001121.

19. Wolf, J., et al. Development of Pandemic Vaccines: ERVEBO Case Study. Vaccines, 2021. 9, DOI: 10.3390/vaccines9030190.

20. Dolzhikova, I.V., et al., Safety and immunogenicity of GamEvac-Combi, a heterologous VSV- and Ad5-vectored Ebola vaccine: An open phase I/II trial in healthy adults in Russia. Human Vaccines & Immunotherapeutics, 2017. 13(3): p. 613–620.

21. Li, Y., et al., Establishing China’s National Standard for the Recombinant Adenovirus Type 5 Vector–Based Ebola Vaccine (Ad5-EBOV) Virus Titer. Human Gene Therapy Clinical Development, 2018. 29(4): p. 226–232.

22. Malik, S. and Y. Waheed, Tracing down the updates on Ebola virus surges: An update on anti-ebola therapeutic strategies. 2023. 11(3): p. 216–225.

23. Lee, J.S., et al., Anti-Ebola therapy for patients with Ebola virus disease: a systematic review. BMC Infectious Diseases, 2019. 19(1): p. 376.

24. Taki, E., et al., Ebanga™: The most recent FDA-approved drug for treating Ebola. Frontiers in Pharmacology, 2023. 14.

25. Saxena, D., et al., Atoltivimab/maftivimab/odesivimab (Inmazeb) combination to treat infection caused by Zaire ebolavirus. Drugs of Today, 2021. 57: p. 483.

26. Tchesnokov, E.P., et al. Mechanism of Inhibition of Ebola Virus RNA-Dependent RNA Polymerase by Remdesivir. Viruses, 2019. 11, DOI: 10.3390/v11040326.

27. Sun, D., et al., Why 90% of clinical drug development fails and how to improve it? Acta Pharmaceutica Sinica B, 2022. 12.

28. Vlachakis, D., Genetic and structural analyses of ssRNA viruses pave the way for the discovery of novel antiviral pharmacological targets. Molecular Omics, 2021. 17(3): p. 357–364.

29. Papageorgiou, L., et al., An updated evolutionary study of Flaviviridae NS3 helicase and NS5 RNA-dependent RNA polymerase reveals novel invariable motifs as potential pharmacological targets. Molecular BioSystems, 2016. 12(7): p. 2080–2093.

30. Dara, S., et al., Machine Learning in Drug Discovery: A Review. Artificial Intelligence Review, 2022. 55(3): p. 1947–1999.

31. Li, W., et al., A Machine Learning-Based Predictive Model for Predicting Lymph Node Metastasis in Patients With Ewing’s Sarcoma. Frontiers in Medicine, 2022. 9.

32. Dong, S., et al., Development and Validation of a Predictive Model to Evaluate the Risk of Bone Metastasis in Kidney Cancer. Frontiers in Oncology, 2021. 11: p. 731905.

33. Davies, M., et al., ChEMBL web services: streamlining access to drug discovery data and utilities. Nucleic Acids Research, 2015. 43(W1): p. W612–W620.

34. Papadatos, G., et al., SureChEMBL: a large-scale, chemically annotated patent document database. Nucleic Acids Research, 2016. 44(D1): p. D1220–D1228.

35. Kanehisa, M.G.S. and S. Goto, KEGG: kyoto Encyclopedia of Genes and Genomes. Nucleic acids research, 2000. 28: p. 27–30.

36. Pickett, B.E., et al., ViPR: an open bioinformatics database and analysis resource for virology research. Nucleic Acids Research, 2012. 40(D1): p. D593–D598.

37. Kim, S., et al., PubChem in 2021: new data content and improved web interfaces. Nucleic Acids Research, 2021. 49(D1): p. D1388–D1395.

38. Polanski, J. and J. Gasteiger, Computer Representation of Chemical Compounds. 2017. p. 1997–2039.

39. Silva, Y., I. Almeida, and M. Queiroz, SQL: From Traditional Databases to Big Data. 2016. 413-418.

40. Landrum, G., RDKit. 2010. https://www.rdkit.org.

41. Goodfellow, I., et al., Generative Adversarial Networks. Advances in Neural Information Processing Systems, 2014. 3.

42. Kingma, D.P. and M. Welling, An Introduction to Variational Autoencoders. ArXiv, 2019. abs/1906.02691.

43. Abadi, M., et al., TensorFlow: A system for large-scale machine learning. 2016.

44. Hodson, T.O., Root-mean-square error (RMSE) or mean absolute error (MAE): When to use them or not. Geoscientific Model Development, 2022. 15: p. 5481–5487.

45. DiPietro, R. and G. Hager, Deep learning: RNNs and LSTM. 2020. p. 503–519.

46. Akhmetshin, T., et al., HyFactor: Hydrogen-count labelled graph-based defactorization Autoencoder. 2021.

47. 2022.02 Chemical Computing Group ULC, -.S.S.W., Montreal, QC H3A 2R7, Canada, Molecular Operating Environment (MOE). 2024. https://www.chemcomp.com/en/Products.htm.

48. Szymański, P., M. Markowicz, and E. Mikiciuk-Olasik *Adaptation of High-Throughput Screening in Drug Discovery—Toxicological Screening Tests*. International Journal of Molecular Sciences, 2012. 13, 427–452 DOI: 10.3390/ijms13010427.

49. Giordano, D., et al. Drug Design by Pharmacophore and Virtual Screening Approach. Pharmaceuticals, 2022. 15, DOI: 10.3390/ph15050646.

50. Prasanna, S. and R. Doerksen, Topological polar surface area: A useful descriptor in 2D-QSAR. Current medicinal chemistry, 2009. 16: p. 21–41.

51. Leung, D.W., et al., Ebolavirus VP35 is a multifunctional virulence factor. Virulence, 2010. 1(6): p. 526–531.

52. Yuan, B., et al., Structure of the Ebola virus polymerase complex. Nature, 2022. 610(7931): p. 394-401.

53. Kalinowsky, L., et al., A Diverse Benchmark Based on 3D Matched Molecular Pairs for Validating Scoring Functions. ACS Omega, 2018. 3(5): p. 5704–5714.

54. Leon, M., et al., Comparing SMILES and SELFIES tokenization for enhanced chemical language modeling. Scientific Reports, 2024. 14(1): p. 25016.

55. van Tilborg, D. and F. Grisoni, Traversing chemical space with active deep learning for low-data drug discovery. Nature Computational Science, 2024. 4(10): p. 786–796.

